# The Latency of Spontaneous Eye Blinks Marks Relevant Visual and Auditory Information Processing

**DOI:** 10.1101/2020.07.23.217547

**Authors:** Supriya Murali, Barbara Händel

## Abstract

Eye blinks are influenced by several external sensory and internal cognitive factors. However, neither the precise temporal effects of these factors on blinking nor how their timing compares between modalities is known. Our aim was to understand the influence of sensory input vs task-relevant information on blinks in the visual and auditory domain.

Using a visual and an auditory temporal judgement task, we found that blinks were suppressed during stimulus presentation in both domains and the overall input length had a significant positive relationship with blink latency i.e. the end of blink suppression. Indeed, the effect of sensory input duration on blink latency was not significantly different between visual and auditory stimuli. The precise timing of blink latency was further modulated by the duration of the task relevant input, which was independent of the overall length of sensory input. The influence of task related changes embedded in the overall stimulation suggests an additional influence of top-down processes on blink timing. Intriguingly, embedded changes as short as 40 ms in the auditory domain and 100 ms in the visual domain are reflected in blink latency differences. Importantly, we could show that task accuracy and motor response was not the driving factor of blink modulation.

Our results show a sensory domain independent modulation of blink latencies introduced by changes in the length of task-relevant information. Therefore, blinks not only mark the time of sensory input but also can act as precise indicator for periods of cognitive processing and attention.

## 1 Introduction

Humans generate about 12 spontaneous eye blinks in a minute (Fatt and Weissman 1992). While one main function is to moisten the corneal tear film, the frequency of their occurrence is much higher than what would be required for that purpose (Zametkin, Stevens and Pittman 1979). Three factors have been suggested to additionally influence the blink rate: sensory input processing, the internal mental state and motor output (Bonneh, Adini and Polat 2016, Cong, et al. 2010, Hoppe, Helfmann and Rothkopf 2018, Oh, Jeong and Jeong 2012, Siegle, Ichikawa and Steinhauer 2008, Wascher, et al. 2015)

Interestingly, not only the rate of blinks is affected by these factors, but also the blink timing. Instead of being randomly executed during a task, blinks follow a specific decrease-increase-baseline pattern, which, Bonneh, Adini, & Polat (2016), refer to as an “oculomotor inhibition mechanism”. This pattern is characterized by decreasing or suppressing one’s blinks during stimulus or information presentation and increasing it during or after the offset of that information (Hoppe, Helfmann and Rothkopf 2018, Oh, Jeong and Jeong 2012, Ohdra 1995, Siegle, Ichikawa and Steinhauer 2008). The purpose of reduced blinking, specifically during visual stimulus presentation, could be simply to not lose any incoming information, which is also a probable reason why visual tasks tend to have a lower blink rate (Goldstein, et al. 1985). However, the decrease is not specific to the visual modality but has been observed during auditory tasks as well (Bauer, et al. 1985, Kobald, et al. 2019) and the overall duration of a stimulus has been found to increase the duration of blink suppression in both domains (Bauer, et al. 1985, Oh, Jeong and Jeong 2012). This suggests a domain independent mechanism. A study by Hoppe, Helfmann, & Rothkopf (2018) even showed that a mere expectation of a visual stimulus can have an influence on one’s blinking which would mean that sensory input is not the driving factor of blink modulation, but that more general cognitive processes influence blinking. In fact, it has been argued that the decrease occurs during periods of attention allocation and that the increase represents the point when all information processing has been completed (Wascher, et al. 2015). This fits to the finding that the suppression of blinks is stronger for more difficult tasks (Oh, Jeong and Jeong 2012).

Apart from the factors pertaining to the stimulus or task itself, blinks have also been shown to be influenced by the motor response. A study by Colzato, van Wouwe, & Hommel (2007) showed that blink rates can reflect the strength of visuomotor binding between a task relevant visual stimulus and the response involving a left and right key press. It has further been shown that blinks are modulated around the motor output (Bauer, et al. 1985, Goldstein, Bauer and Stern 1992, Oh, Han, et al. 2012) and can be entrained to the motor response even when participants engage in self-paced rhythmic finger tapping without external sensory cues (Cong, et al. 2010). Nevertheless, motor aspects seem not to be the sole influence on blink occurrence since blinks have been shown to be modulated even during no-go trials (Kobald, et al. 2019, Wascher, et al. 2015).

We wanted to understand and dissociate the influences of motor response, sensory input and cognitive processes on blink behavior and rigorously compare it between sensory domains. To this end, we independently manipulated task irrelevant and task relevant sensory input during a comparable auditory and visual task, and considered the motor output as well as more cognitive influences related to the task performance. We specifically focused on the temporal effects on blinking behavior by looking at the blink latencies, a measure which has been used previously to detect influences on blinking (Doughty 2001, Fogarty and Stern 1989, Goldstein, et al. 1985, Goldstein, Bauer and Stern 1992, Kobald, et al. 2019).

We conducted a visual and auditory temporal judgement task in which subjects had to judge if two stimuli appeared simultaneously. To secure comparable performance between the two modalities smaller time differences between the two stimuli were used in the auditory task, since temporal processing has been shown to be better for auditory input (Kanabus, et al. 2002). Also, since low level stimulus features can effect blinking (Bonneh, Adini and Polat 2016), we conducted the experiments in complete darkness. Our results showed that blinks were suppressed similarly in the two domains and that the length of this suppression correlated with the duration of the task-relevant information periods independent of the overall stimulus length or the motor response. Hence, our findings suggest a temporally precise domain general top-down modulation of eye blinks.

## 2 Methods

Eighteen subjects (4 males) aged between 18-35 years participated in the study. All participants gave their written informed consent and received either payment or study credit for their participation. The study was approved by the local ethics committee and was in line with the European general data protection regulations (DSVGO).

Two subjects were excluded for the visual condition because of an overall response accuracy below 10% and a mean blink rate of below 1 blink per minute respectively and one subject was excluded in the auditory condition for a low blink rate. Six additional subjects were excluded for the analysis pertaining to the ISI and Stimulus ON-time (i.e. duration until both stimuli were turned off) in the auditory condition, due to incomplete data recordings.

The mobile SMI Eye tracker glasses (ETG 2W Analysis Pro-120Hz) was used to record eye data and responses were given via two response buttons connected to a response box (model: K-RB1-4; The Black Box ToolKit Ltd, UK), which was connected to a Dell Precision (M6700) laptop via a USB cable. The experiment and the analysis were coded in MATLAB 2012 and 2015a using the Psychtoolbox extensions (Brainard 1997, Kleiner , Brainard and Pelli 2007, Pelli 1997). All data streams were recorded using the Lab Streaming Layer, LSL (https://github.com/sccn/labstreaminglayer) along with LabRecorder (version 1.12b). The experiment was conducted in a completely dark room.

### 2.1 Visual Condition

The visual stimulus was presented using three red light emitting diodes (LEDs) with a diameter of 4mm, placed on a metal ruler using magnets. The LEDs were placed in front of the subject at a distance of 50 cm at eye level in a completely dark room (modified EEG booth). The central LED was only turned on at the start of the experiment and was switched off during the trials. The other two LEDs were placed on either hemifield, each at 11 degrees from the central fixation. During each trial, the two stimuli (LEDs) turned on at the same or at different times with inter-stimulus intervals (ISI) ranging from 0 to 0.2s in steps of 0.02s. The 0 ISI indicates that both stimuli turn on simultaneously. The order of the stimuli (for ISI>0) was randomized within subjects. After their onset, the two stimuli remained on for 0.4, 0.5 or 0.6s, which is referred to as the stimulus ON-time (Duration both stimuli were on until the time both were turned off). Also this time period was randomly assigned per trial. After this, both LEDs were turned off. To further add to the unpredictability of the next trial, we jittered the next stimulus onset by randomly adding an inter-trial interval between 0.5 and 0.6s (see Figure 1a). The participants had to indicate if the two lights turned on at the same or at different times by pressing either a left or a right button (randomly assigned for each subject) as fast as possible at any time during the trial. There were two sessions of 506 trials each, adding to a total of 1012 trials. We included one long break in between the sessions and two short breaks within each session to avoid excessive fatigue, which can be introduced by sitting in complete darkness.

**Figure 1.**
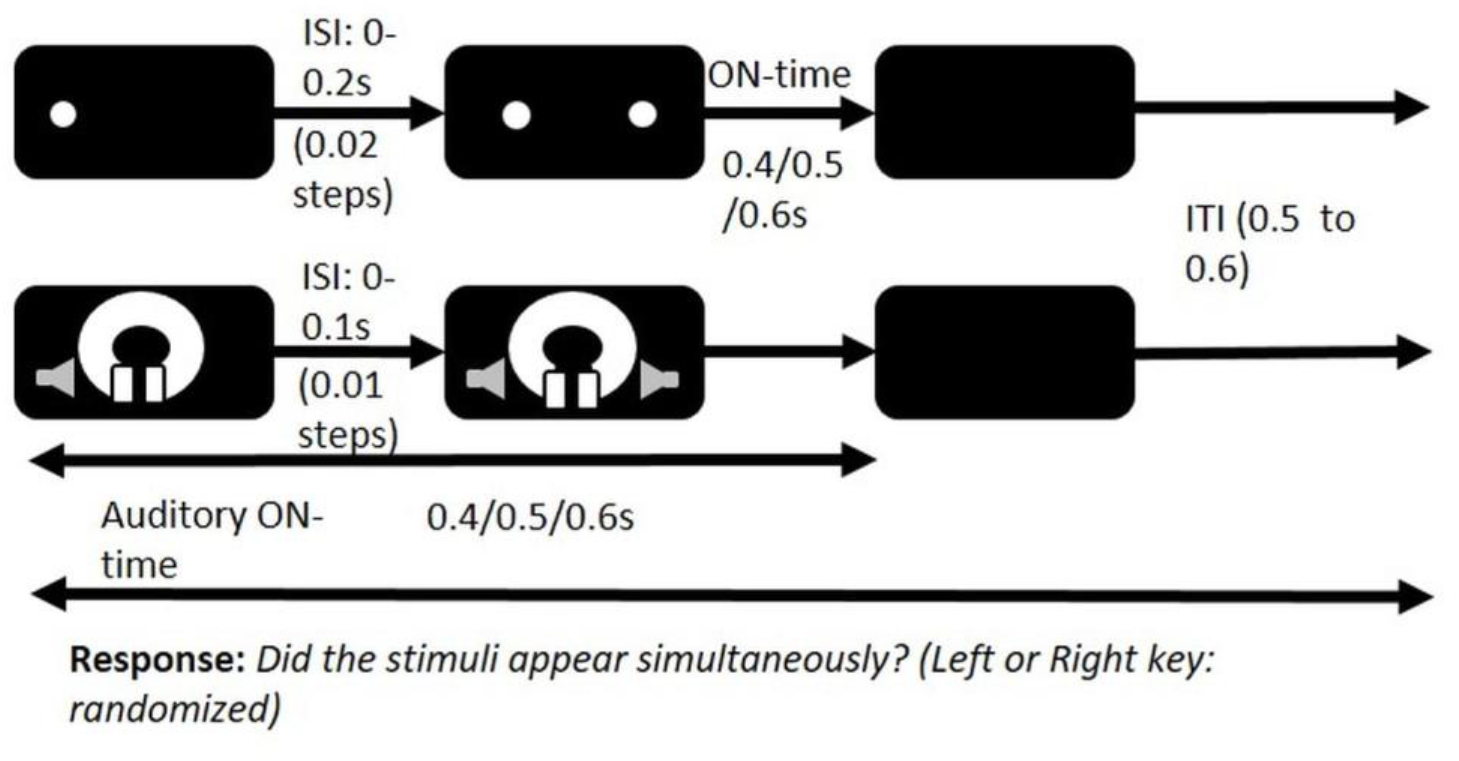
Stimulus and Task during the visual and the auditory conditions

### 2.2 Auditory Condition

The auditory condition was similar in structure and task but now the stimulus consisted of two sound clicks (500 Hz) that were presented in each ear using the Sennheiser PC3 headset along with the Steinberg UR22mk II external soundcard, again in complete darkness. The procedure and task was similar to the visual condition (see Figure 1b), with three differences: 1) the ISIs did not have an influence on the trial length i.e. the stimulus ON-time refers to the duration at least one stimulus was on until the time both were turned off) 2) The ISIs ranged from 0 to 0.1s in steps of 0.01s. The lower ISI range for auditory was used since auditory discrimination thresholds have been found to be lower (see Introduction) 3) There was only one session of 506 trials, whereas the visual condition had two sessions. The reason for using two sessions in the visual condition was to obtain enough blinks, since blink rates are generally lower during visual tasks (see Introduction).

### 2.3 Stimulus Timing Test

Before we started with data collection, we tested our stimulus to make sure that the timing was precise. This was done using a photodiode for the visual and an audio output recorder that measured the audio output directly from the audio jack.

## 3 Analysis

### 3.1 Blink Detection

We computed a blink detection algorithm based on the pupil radius data. The z value of the radius data was calculated and a potential blink was considered if either 1) both radii values were missing or 2) if either the left or right eye data was missing and the other eye had a z-value below a threshold (−1, −2 or −3, set individually because of considerable difference in signal to noise ratio in the data).

The onset and the offset of the blink were computed based on the last point, for either eye, where the z-values was still above the threshold before the blink, or the first z-value that reached above the threshold after the blink. Blinks with less than 0.016s between them were combined and those with durations lower than 0.05s or higher than 0.5s were discarded.

### 3.2 Blink rates (time resolved)

The plotted time series consists of 0.1s non-overlapping time windows. We calculated the normalized mean number of blinks in each time window by dividing the mean number of blinks in that time window by the mean number of blinks in all time windows.

### 3.3 Blink latency

We also looked at three factors that could modulate the timing of blink occurrence: the stimulus ON-time (time until both stimuli were turned off), the ISI and the reaction time. We took the mean blink latency as the dependent measure (discussed in the next part).

For the factor ON-time, the blink latency was calculated as the difference between the first blink onset and the onset of the second stimulus i.e. discarding the influence of the ISI period. To account for the possibility that merely taking the overall average for each ON-time might lead to differences due to different number of blinks for each ISI within each ON-time, we first calculated the mean over each ISI within the ON-time and then averaged over these ISI specific means. This was followed by a two-factor ANOVA with factors ON-time (0.4, 0.5, 0.6) and Modality (Visual vs. Auditory) and the mean Blink latency as the dependent variable.

For the factor ISI, a regression analysis was conducted. The dependent variable, the blink latency, was taken as the difference between the first blink onset and the trial onset. We followed the same procedure as describe above for the ON-time, by calculating the mean over each ISI within each ON-time condition first and then calculated an overall mean.

Lastly, for the reaction time, we first calculated the mean blink latency from the trial onset and the mean reaction time (also from the trial onset) for each subject. A regression analysis with the reaction time as the predictor and blink latency as the dependent variable was then conducted.

## 4 Results

### 4.1 Overall Performance (accuracy and reaction time)

The overall mean accuracy and reaction time for the visual condition was 48.5% (*SD* = 15.4%) and 0.56s (*SD* = 0.22s) respectively; and for the auditory condition 63.8% (*SD* = 22.5%) and 0.5s (*SD* = 0.14s) respectively. For both the visual and the auditory conditions, the accuracy was highest for 0 ISI since it was easy to perceive the LEDs as lighting up or the tones appearing simultaneously. As further expected, the accuracy increased with increasing ISIs (for ISIs above 0) (Figure 2a and b). Regression analysis showed a significant linear relationship between accuracy and ISI (calculated from 0.02s for the visual and from 0.01s for the auditory condition) for both the visual (*R*^*2*^ = 0.98, *b* = 0.02, *F* (1,8) = 361.1, *p* < .001) and the auditory (*R*^*2*^ = 0.90, *b* = 0.4, *F* (1,8) = 72.4, *p* < .001) condition.

**Figure 2.**
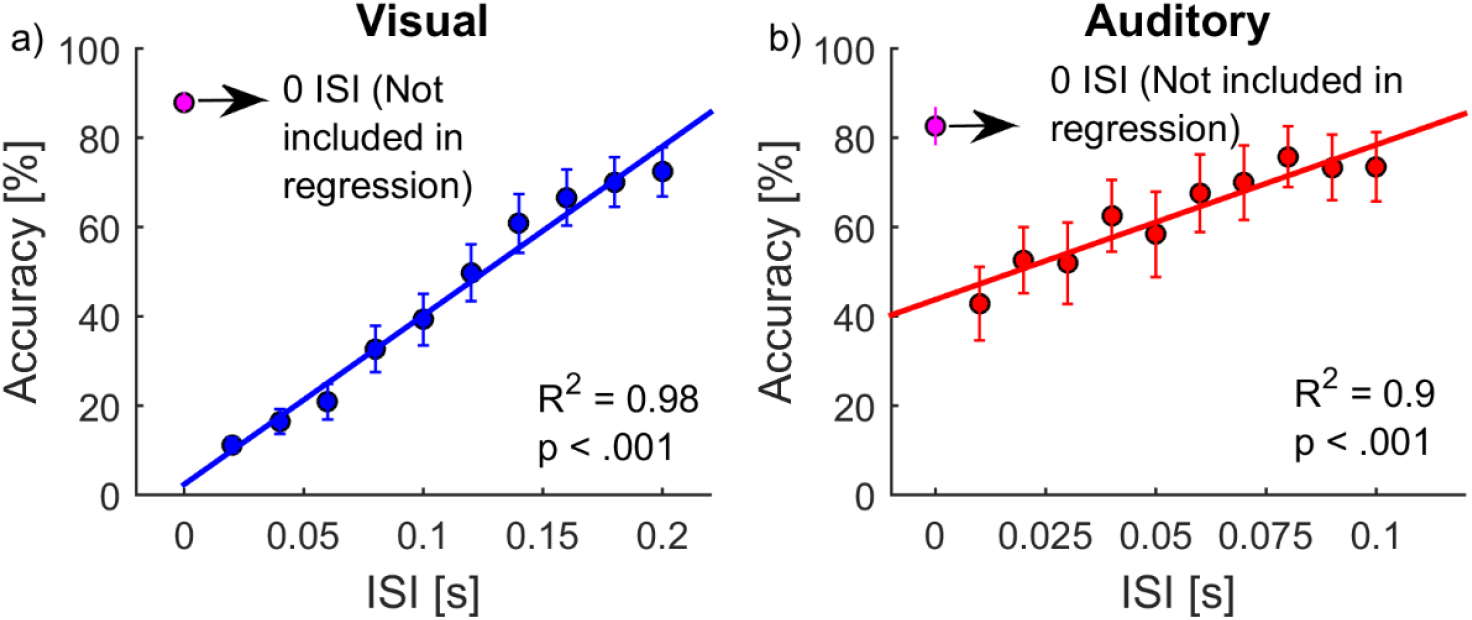
Interstimulus interval (ISI) vs. Accuracy. a) During the visual condition b) During the auditory condition. Error bars represent standard errors.

### 4.2 General Blink Parameters (overall rate and duration)

The mean blink rate during the visual condition was 7.8 per minute (*SE* = 2.02) and for the auditory it was 11.5 per minute (*SE* = 2.7), as shown in Figure 3. T-test showed a significant difference between the blink rates (*t* (14) = 2.6, *p* = .02). The mean duration of blinks was 0.14s (*SE* = 0.01 for the visual and 0.15s (SE = 0.02) for the auditory conditions.

**Figure 3.**
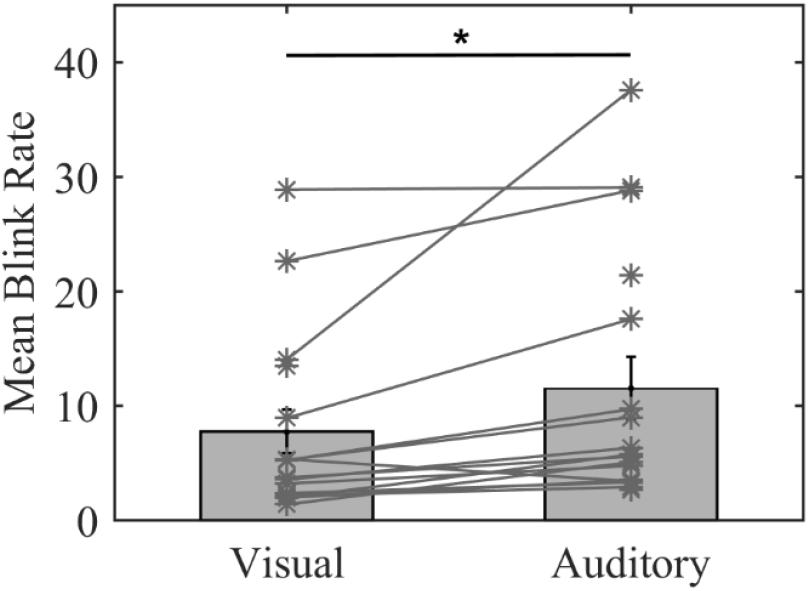
Mean blink rates during the visual and the auditory conditions. Error bars represent standard errors.

### 4.3 Time Resolved Blink Rates

Special interest was placed on the modulation of the blink rate over time within a trial. Specifically, we looked at the mean blink rate from 0.2s before the trial onset to 2s after the trial onset for both the modalities (Figure 4). The purpose was to see if we can replicate the previously described modulation in the visual domain and if it similarly occurs in the auditory domain. The graphs show a similar modulation in both modalities with an increase in blink rate starting about 0.3s lasting until 1.2s after the trial onset.

**Figure 4.**
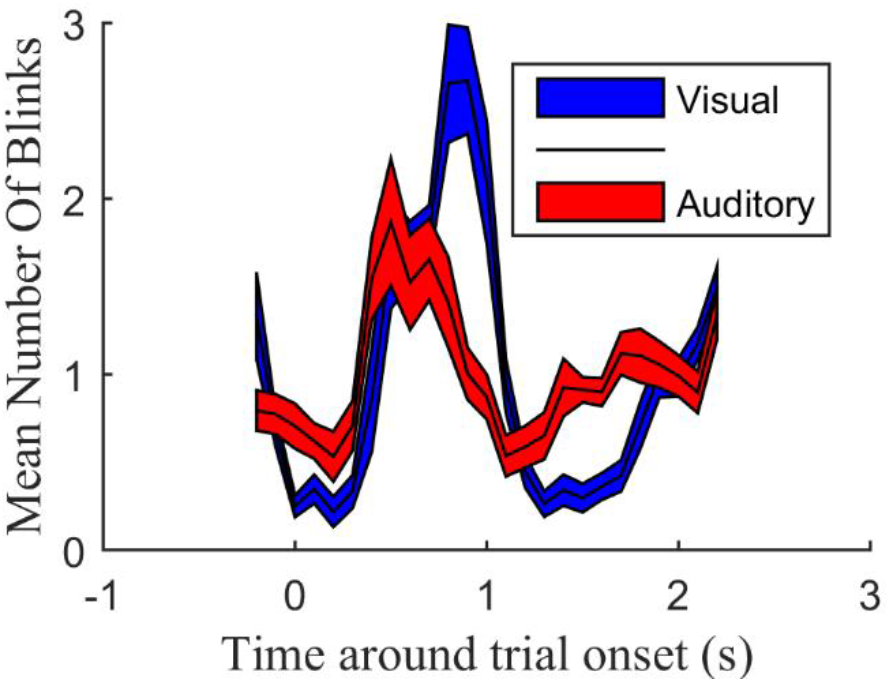
Normalized mean number of blinks during the visual and the auditory condition. The x-axis represents timing with 0.1s non-overlapping windows. The y-axis represents the normalized mean number of blinks in each window over all subjects. The colored region represents the standard error.

### 4.4 Blink Latency

In order to understand what influences the timing of this modulation, we looked at the latency of the first blink during each trial. Specifically, we investigated the possible influence of the task-independent sensory input defined by the ON-time (Time until the stimuli were turned off), the task-related input (ISI) and the timing of the motor response reflected in the reaction time.

#### 4.4.1 Factor 1-Stimulus ON-time (time until both stimuli were turned off): (Task-independent sensory input)

Figure 5 shows the mean blink latency for the three ON-times in the visual (blue bars) and auditory (red bars) conditions. A two factor repeated measures ANOVAs with factors Modality and ON-time, revealed only a significant effect of ON-time on the blink latency (*F* (2,61) = 7.8, *p* = .001) and no significance for modality (F < 1).

**Figure 5.**
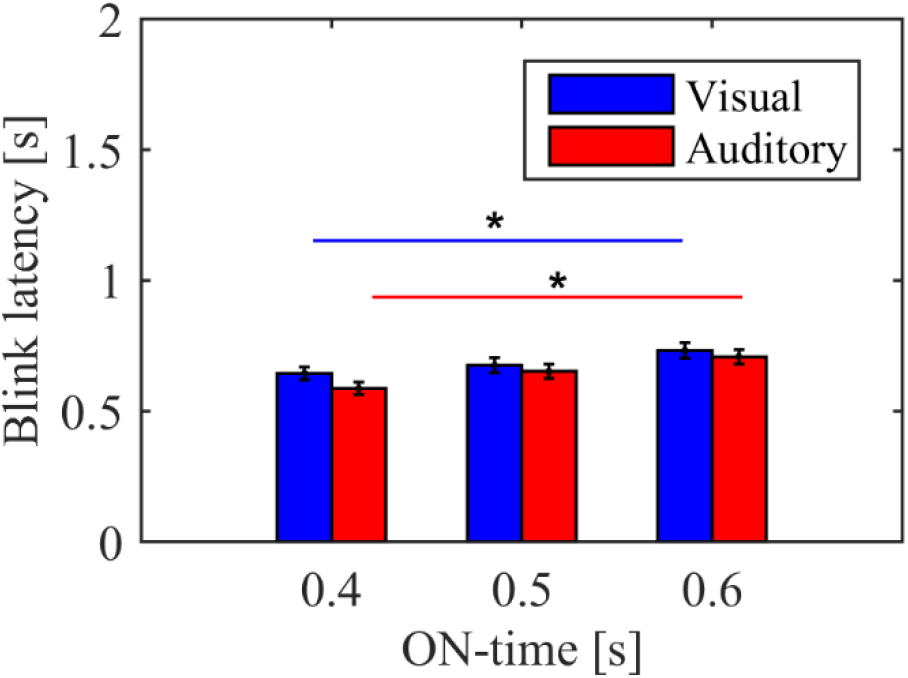
Comparison of blink latencies between the three ON-times (time until both stimuli were turned off) for the visual (blue) and the auditory (red) condition. The error bars represent standard error. Asterix marks significant differences at the level of 0.05 alpha based on a post-hoc Ttest.

PostHoc t-tests revealed, for the visual condition, a significant difference between 0.4s (*M* = .6, *SE* = .02) and 0.6s (*M* = .73, *SE* = .03) ON-time (*t* (15) = 3.2, *p* = .005); but not the between the 0.4 and 0.5s ON-time (*t* (15) = 1.8, *p* = .09) and the 0.5s (*M* = .68, *SE* = .03) and 0.6s ON-time (*t* (15) = 2.1, *p* = .05). Similarly, for the auditory condition, the tests showed a significant difference between 0.4s (*M* = .6, *SE* = .03) and 0.6s (*M* = .7, *SE* = .03) ON-time (*t* (10) = 3.6, *p* = .004); but not between 0.4s and 0.5s (*M* = .6, SE = .03) ON-time (*t* (10) = 2.02, *p* = .07) and 0.5s and 0.6s ON-time (*t* (10) = 1.5, *p* = .16). Longer stimulus ON-time therefore lead to a longer blink latency.

#### 4.4.2 Factor 2: ISI (Task-dependent input)

Increasing ISIs denoted increasing input durations of task relevant information. However, ISIs also positively correlate with accuracy, which could lead to cognitive influences. We therefore needed to dissociate accuracy effects from input duration effects. As shown in Fig 2, the ISIs showed a positive relationship with accuracy with the exception of the synchronous onset (0 ISI), which showed accuracies comparable to the highest ISI (Figure 2a and b). Hence, if the effect of the ISI is due to the increase in input duration, the synchronous onset would follow a positive correlation with blink latency. However, if it is due to accuracy effects, the 0 and the highest ISIs would have comparable latencies (since they led to similar accuracy).

As a first step, we looked at whether there is an effect of the ISI, without considering the 0 ISI, for the visual and the auditory condition respectively (Figure 6a and b). The regression analysis showed that the ISI significantly predicted the blink latency for both the visual (*R*^*2*^ = .46, *b* = .731, *F* (1,8) = 6.98, *p* =.03) and the auditory (*R*^*2*^ = .41, *b* = .64, F (1,8) = 5.65, *p* = .04) condition. Please note that for the auditory condition, the ISIs did not lead to an overall change in stimulus length.

**Figure 6.**
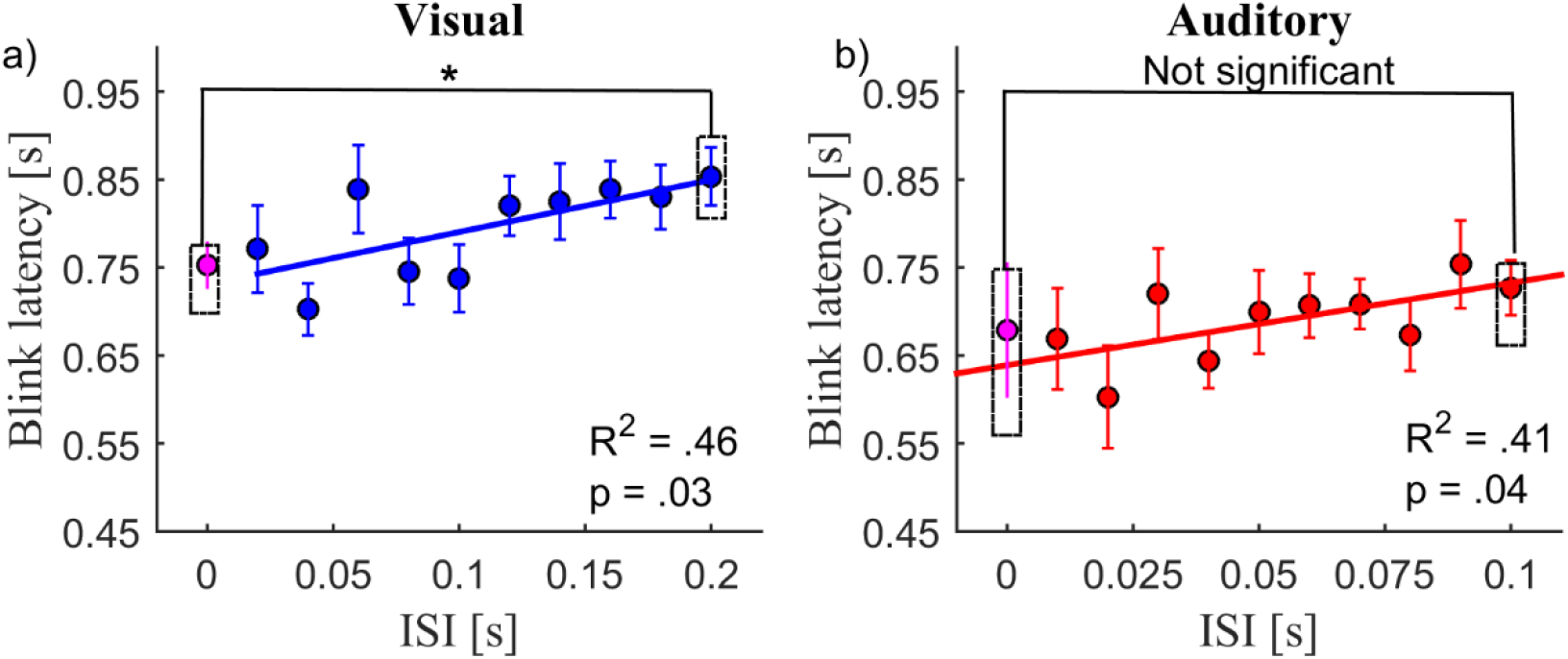
ISI vs Blink latency. Regression analysis did not include the 0 ISI. Ttest (represented in dotted lines) conducted between the 0 ISI and the highest ISI (0.2s for the visual and 0.1s for the auditory). Error bars represent standard errors.

We then compared the 0 ISI with the highest ISI in both conditions (also shown in the same Figure 6a and b). For the visual task (Figure 6a, we found a significant difference (*t* (15) = 3.8, *p* =.002) between the 0 ISI (*M* = .73, *SE* = 0.02) and the 0.2 ISI (*M* = .85, *SE* = 0.035. However, for the auditory condition (Figure 6b), we found no significant difference (*t* (10) = .4, *p* = .705) between the 0 ISI (*M* = .68, *SE* = .09) and the 0.1 ISI (*M* = .72, *SE* = .03).

Additionally, to see how the increase in ISI leads to increase in blink latency, we plotted a psychometric function (Figure 7) based on the data from Figure 6. Specifically, for the visual task, we compared ISIs that were 0.02s upto 0.140s (in steps of 0.02s) apart. For the auditory task we used 0.01 to 0.07 s ISI difference (in steps of 0.01s). For instance, a difference in ISI of 0.04s, would mean that we compare data (blink latency) for the ISI of 0.02 to the one for ISI 0.6, 0.04 to 0.8 and so on untill 0.16 to 0.2. We then calculated how many of these pairs showed an increase in blink latency. This results in a percentage of data points showing a positive change in blink latency dependent on how large the time difference was between the ISIs. Note that, since the number of available data points decrease as we increase the difference in ISI, we refrain from statistics but add a psychometric function for description of the trend. Looking at Figure 7, the function suggests that a change in ISI of 0.1s for visual and 0.04s for auditory leads to an increase in blink latency 80% of the time.

**Figure 7.**
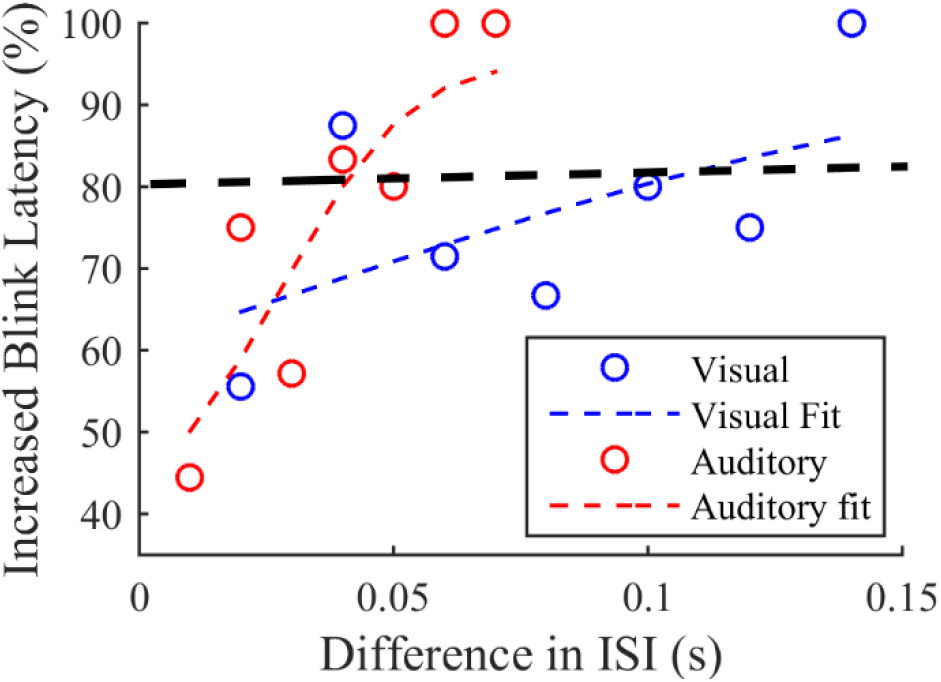
The percentage of cases (including a psychometric fit) in which an increase of ISI of a certain length led to a concurrent increase in blink latency. The x-axis shows the difference between the ISIs that are taken to compare the blink latency (as taken from Figure 6). The y-axis shows the percentage of data points were in an increase in ISI (as specified on the x-axis) led to an increase in blink latency

#### 4.4.3 Factor 3: Reaction time

Figures 8a and b show the relationship between the reaction time and the blink latency. The regression analysis showed no significant influence of the RT in either of the conditions (visual: *R*^*2*^ = .11, *b* = 1.1, *F* (1,14) = 1.65, *p* = .22; auditory: *R*^*2*^ = .11, *b* = 0.46, *F* (1,15) = 1.92, *p* =.2) and additionally shows a reversed slope for the auditory and visual condition.

**Figure 8.**
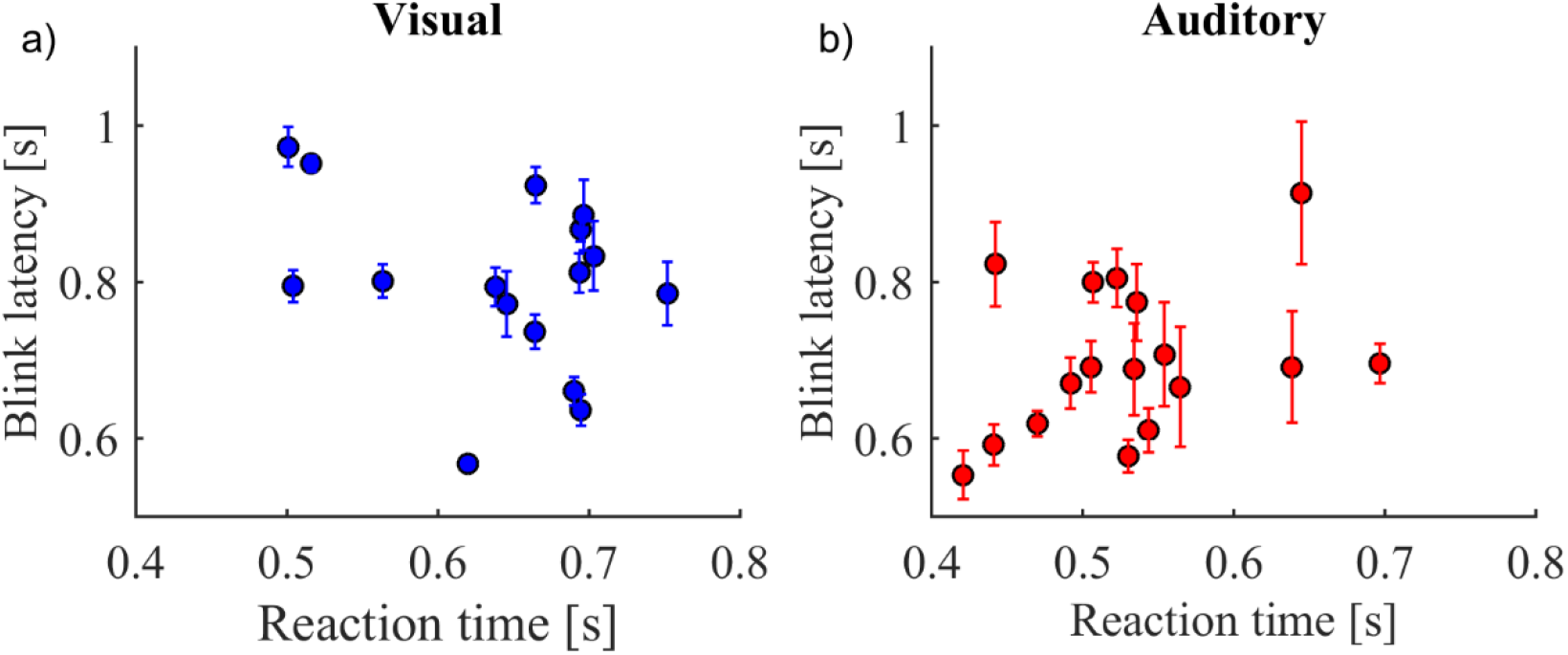
Regression analysis between the Reaction time and Blink Latency the visual (a) and the auditory (b)

## 5 Discussion

Using an auditory and visual simultaneity judgment task, our study shows a temporally precise modulation of blink latency influenced by sensory as well as cognitive factors, similarly in the auditory and visual domain. Specifically, periods involving stimulus presentation were associated with a low blink rate and an increase after its offset. Our aim was to understand what influences this modulation by investigating the duration of the period of reduced blinking during stimulus presentation by looking at the timing of the first blink after the onset of the stimulus, referred to as blink latency. This time can be interpreted as period of blink suppression. We specifically looked at the influence of the following factors on the blink latency: the overall sensory input, the time of the manual response (reaction time), the task specific sensory input (ISI) and indirectly at task accuracy. A rigorous comparison between these different factors in a comparable visual and auditory task is missing to date.

The overall sensory input duration showed the most robust effect, in the sense that, people always tended to wait until the offset of the sensory input before starting to blink. Figure 5 shows that the blink latency was approximately 0.6s for a stimulus ON-time (time until both stimuli were turned off) of 0.4s. Note that, the latency was calculated from the time both stimuli were turned on, which was the beginning of the 0.4s period and the end of this period marks the stimulus offset. Hence, a blink was most likely to occur after the stimulus was turned off. The pattern of reduced blink rate followed by an increase has been reported before in both the visual (Oh, Jeong and Jeong 2012, Siegle, Ichikawa and Steinhauer 2008) as well as the auditory (Oh, Han, et al. 2012, Oh, Jeong and Jeong 2012) domains. It was interpreted as an active suppression followed by its rebound (Hoppe, Helfmann and Rothkopf 2018, Oh, Jeong and Jeong 2012). In fact, it has been suggested that blinks follow the same inhibitory mechanism as other eye movements during the processing of any sensory event (Bonneh, Adini and Polat 2016).

In the visual domain, this suppression could be mediated by the desire to not lose any information; which is also why blink rates are usually lower for visual tasks (Goldstein R. W., 1985). However, similar to other researchers (Oh, Han, et al. 2012, Oh, Jeong and Jeong 2012), we also found the same modulation of blink latency in the auditory domain, which indicates that an interpretation based on visual input processing is insufficient. Indeed, it has been argued that the decrease in blinks is associated with attention to the input, where the subsequent increase in blinks marks the end of the attentional period i.e. when all information processing is complete (Kobald, et al. 2019, Wascher, et al. 2015). The suppression of blinks until its release is possibly a result of attentional allocation to sensory input. In fact, a study by Hoppe, Helfmann, & Rothkopf (2018) showed that even a mere increase in the probability of a visual event was sufficient to lower the probability of blinks, strengthening the idea of a non-sensory driven active top-down influence. Therefore, a precise temporal modulation in blinks through top-down factors could occur independent of the sensory domain and sensory input.

Indeed, our further findings suggest such a domain general attentional effect in blink suppression independent of the overall sensory input duration. We found that task-specific sensory input (ISI), embedded in the overall sensory input, was significantly associated with increasing blink latencies (Figure 6a & b). Subjects still waited on average until the overall stimulation offset, but the length of the task-relevant (to be attended) input added time to the sensory based (bottom-up driven) latency. Please note that we argue here based on the auditory results, since in the visual task, the ISI added to the overall input length (Figure 1). However, since the main effects of the different ON-times (time until both stimuli were turned off) were comparable between the two modalities, this difference might be negligible. Assuming that subjects wait until both LEDs/ tones have been presented before deciding if they turned on simultaneously or not, the ISI increased the duration the relevant input needed to be complete. This would increase the length of attended time within the overall sensory input. Our data (excluding simultaneous onset) showed a strong positive correlation with blink latency. Increased ISI however is also highly correlated with accuracy. Accuracy increases with the increase in ISI (Figure 2a and b), except (not surprisingly) for the 0 ISI which is simple to detect as simultaneous input. This specific case helps to understand that it is not accuracy related effects that drive the blink latency, but indeed the length of task relevant input. Nevertheless, there might still be an influence of task difficulty as shown by (Drew 1951, Goldstein, Bauer and Stern 1992, Veltman and Gaillard 1998) which is masked by the influences of sensory input. We argue against a role of accuracy in explaining our findings in the following paragraph.

While accuracy is similarly high for 0 ISI in both modalities, the time of relevant input however is different for the two modalities. This stems from a basic perceptual difference between the modalities. For visual input, two spatially non-overlapping but temporally overlapping signals can be perceived easily as two separate inputs, whereas this is not the case for a temporally overlapping tone presented to the left and right ear. Two signals that are sufficiently correlated between the ears are fused and interpreted as a single auditory event (Blauert 1938). This is known as binaural fusion and has been widely studied (Broadbent 1955, Leakey 1958, Masakazu 1991). In our task, this fusion forces subjects to wait until the probability of the second tone to occurred is close to zero (which would be after the highest possible ISI period), before being able to decide what answer to give while the answer itself will be very accurate. Therefore, while accuracy for 0 ISI is similar for the auditory and visual task, the time attention needs to be focused on the input differs between modalities. This can explain why the 0 ISI in the auditory condition led to a blink latency that exceeded the one of the highest ISI while the 0 ISI in the visual task has the lowest blink latency. This finding clearly suggests that it is not accuracy related effects but the time of task relevant input (and likely its attention focused on it) which influences blink latency similarly in both sensory domains. Since task relevance is defined by the instruction and independent of the overall sensory input, a top-down influence on the blink latency can therefore be assumed.

Finally, our results show that the motor response or its planning played no part in the modulation of blink latency. Although, motor output can have an independent influence on blinking (Cong, et al. 2010, van Dam and van Ee 2006), our data (Figure 8a & b) showed no correlation between the manual response and blink latencies.

In summary, while a decrease in blinks was previously associated with sensory input in the visual domain, we find the same modulation during purely auditory input. Additionally, our results show that minute changes of task relevant information length, even if embedded in ongoing sensory stimulation, modulate blinking behavior similarly in the auditory and visual domain (Figure 7a & b). Our study therefore highlights domain general top-down influences that can precisely modulate the timing of blinking, mapping small temporal changes in sensory-attentional demands.

## Funding

This project has been funded by the European Research Council (grant number 677819 awarded to B. Händel).

## Notes

### Competing Interest Statement

The authors have declared no competing interest.

